# PoMeLo: a systematic computational approach to predicting metabolic loss in pathogen genomes

**DOI:** 10.1101/2023.08.01.551502

**Authors:** Abigail Leigh Glascock, Eric Waltari, Gytis Dudas, Joan Wong, Vida Ahyong

## Abstract

**Background:** Genome streamlining, the process by which genomes become smaller and encode fewer genes over time, is a common phenomenon among pathogenic bacteria. This reduction is driven by selection for faster replication and minimized energy expenditure in a nutrient-rich environment. As pathogens evolve to become more reliant on the host, metabolic genes and resulting capabilities are lost in favor of siphoning metabolites from the host. Characterizing genome streamlining, gene loss, and pathway degradation can be useful in assessing pathogen metabolic dependency on host metabolism and identifying potential targets for host-directed therapeutics.

**Results:** PoMeLo (**P**redictor **o**f **Me**tabolic **Lo**ss) is a novel evolutionary genomics-guided computational approach for identifying metabolic gaps in the genomes of pathogenic bacteria. PoMeLo leverages a centralized public database of high-quality genomes and annotations and allows the user to compare an unlimited number of genomes across individual genes and pathways. PoMeLo runs locally using user-friendly prompts in a matter of minutes and generates tabular and visual outputs for users to compare predicted metabolic capacity between groups of bacteria and individual species. Each pathway is assigned a Predicted Metabolic Loss (PML) score to assess the magnitude of genome streamlining. Optionally, PoMeLo places results in an evolutionary context by including phylogenetic relationships in visual outputs. It can also initially compute phylogenetically-weighted mean genome sizes to identify genome streamlining events. Here we describe PoMeLo and demonstrate its use in identifying metabolic gaps in genomes of pathogenic *Treponema* species.

**Conclusions:** PoMeLo represents an advance over existing methods for identifying metabolic gaps in genomic data, allowing comparison across large numbers of genomes and placing the resulting data in a phylogenetic context. PoMeLo is freely available for academic and nonacademic use at https://github.com/czbiohub-sf/pomelo.

## Background

All obligate bacterial pathogens begin, evolutionarily, as free-living organisms. As they evolve to become more reliant on their hosts, metabolic genes and resulting capabilities are lost in favor of siphoning metabolites from the host [1, 2]. Over time, the pathogen can become completely reliant upon the host for one or more essential metabolites. Reductive evolution, or genome streamlining, is the process by which microorganism genomes become smaller and less complex over time and can be particularly pronounced during evolution towards symbioses [3, 4]. It is thought to be driven by a preference for faster replication and minimized energy expenditure, and is often hastened in an environment in which metabolites are freely available (e.g. a host). Consequently, genome streamlining has been observed across many distinct lineages of bacterial pathogens including *Treponema pallidum*, the causative agent of syphilis, and *Mycoplasma pneumoniae*, a common cause of bacterial pneumonia [5, 6].

To computationally predict metabolic capabilities of different organisms, genome annotation and reconstruction of metabolic pathways is required. The widely used RAST and KEGG servers allow users to perform basic annotations and comparisons on a small number of genomes [7, 8]. Here we describe PoMeLo (**P**redictor **o**f **Me**tabolic **Lo**ss), a novel computational approach to analyze metabolic loss in bacterial genomes. Given two sets of genome groups – target and non-target species – PoMeLo systematically characterizes and compares the metabolic capabilities of any number of bacterial genomes via a user-friendly interface. An optional phylogenetically-weighted mean genome size analysis allows users to identify evolutionary branch points at which genome streamlining has potentially taken place, to inform their initial selection of target and nontarget genomes. PoMeLo performs comparative analyses across individual genes as well as full pathways, and it summarizes the results at the group (target/non-target) and species levels. Importantly, a Predicted Metabolic Loss (PML) score is computed for each pathway to illustrate the magnitude of gene loss across all individual pathways. PoMeLo leverages a large public microbial genome repository, in which all genomes and annotations have been generated in the same manner, ensuring consistency across multiple comparisons. The tool runs locally in a matter of minutes, generating tabular and visual outputs that allow users to compare predicted metabolic functionality between groups of genomes and individual species.

As a demonstration, we use PoMeLo to perform an analysis of genomes within the genus *Treponema*. Both *T. pallidum* and *T. paraluiscuniculi*, the causative agents of syphilis in humans and rabbits, respectively, show strong evidence for having undergone extensive genome reduction [5, 9], making them good candidates for interrogating metabolic pathway changes in comparison to related bacterial groups. These organisms have an average genome size of 1.1 Mb, while genomes of close relatives such as *T. phagedenis* (2.9–3.8 Mb) are all greater than 2.5 Mb in size. Thus, we selected *T. pallidum* and *T. paraluiscuniculi* as the target group and *T. phagedenis* as the non-target group in our analysis. We show that PoMeLo facilitates high-throughput, rapid analysis of metabolic loss in bacterial genomes, providing a Predicted Metabolic Loss score for each metabolic pathway and placing the results in evolutionary context.

## Implementation

### Running PoMeLo

Before running PoMeLo, the user must have an internet connection, download the `PML_script.R` script and the `mapping_GO_to_ecgene_and_ ecpathway_toPATRIC.tab` mapping file (Supplementary Table 1) from the GitHub repository (https://github.com/czbiohub-sf/pomelo), install the required R packages (Supplementary Table 2), and load them using the `library()` function in R. PoMeLo accepts as input two lists of genomes designated as “target” and “non-target.” The functional annotations associated with the input genomes are used to predict the metabolic potential of pathogens of interest in comparison to their free-living relatives.

### Genome Selection

Two lists of target and non-target genomes must be generated via the Bacterial and Viral Bioinformatics Resource Center (BV-BRC) [10] and downloaded from the their website (https://www.bv-brc.org/), formatted as comma-separated values (CSV) files.

Target genomes should be reduced in size and/or gene content in comparison to nontarget genomes, so that differences in metabolic potential are more easily discerned (see Optional Genome Loss/Gain Analysis section). It may be necessary to include non-target genomes that are more evolutionarily distant from the target group to observe genome reduction and streamlining, especially in cases where significant genome reduction has occurred across the entire genus (e.g. *Mycoplasma*). Users can navigate to their organisms of interest via either the search bar on the home page or the ‘`Organisms`’ tab at the top of the page, by selecting ‘`All Bacteria`’ and navigating through the resulting page. It may be helpful for users to explore the ‘`Phylogeny`’ tab to view which organisms have representative genomes available within the BV-BRC database and how they are related to one another.

To generate the list of target genomes, users can click on the ‘`Genomes`’ tab and filter the results by clicking on the ‘`FILTERS`’ icon. To ensure that pathway annotations and metabolic predictions are accurate, high quality genomes should be used for this analysis. The authors suggest filtering by ‘`Genome Quality`’ for ‘`Good`’ quality genomes and filtering by ‘`Genome Status`’ for ‘`Complete`’ genomes. After filtering the genome list, the user can select genomes for download by checking the boxes to the left of the genome name. Once the target genomes have been selected, the user can click the ‘`DOWNLOAD`’ icon and select the ‘`CSV`’ option. This will trigger an automatic download of the genomes list in CSV format. This process should then be repeated to select and download the list of non-target genomes.

### Optional Phylogenetic Analysis

As a part of the PoMeLo GitHub repository, we have provided an additional script (`PML_PICanalysis.Rmd`) that takes as input a list of genomes of interest along with a user-supplied phylogeny of the genomes in Newick format (NWK). The script estimates the genome size gain or loss across the phylogeny by computing phylogenetically-weighted mean genome sizes. The phylogeny can be constructed using the BV-BRC resource by the following steps: 1) select one representative genome for each species in the ‘`Genomes`’ tab, 2) create a genome group by clicking on the ‘`GROUP`’ icon, 3) navigate to the ‘`Tools & Services`’ dropdown menu and select ‘`Bacterial Genome Tree`’ under the ‘`Phylogenomics`’ header, and 4) build a tree using the previously defined group. The phylogenetically-weighted mean genome size analysis takes shared evolutionary history into account and follows the first steps of phylogenetically independent contrasts (PICs) but skips the final standardization by the variance [11–13].

At each internal node of the tree, the algorithm computes phylogenetically-weighted mean genome sizes by summing the actual values (for external tips, i.e. actual genomes) or computed values (for internal nodes) of its descendants and dividing the total by the number of descendant branches of that node. These estimates are propagated from the external tips to the root of the tree (Supplementary Figure 1). The algorithm then estimates the percentage change in genome size along each branch. This allows the user to visualize the genome size changes for each species and each progressively distant clade in the tree. Using this approach, users can easily pinpoint branches in evolutionary history where genome size has been reduced and estimate the magnitude of that loss. In conjunction with biological knowledge of the organism, its pathogenicity, and localization within the host (e.g. extracellular, vacuolar, cytoplasmic), users can more easily distinguish which species to select for target versus non-target groups. For accurate prediction of metabolic loss in pathogen genomes, it is important for the target group genomes to have undergone sufficient gene loss.

Optionally, the two files generated for genome size analysis can also be used to incorporate phylogenetic information to the visual outputs of PoMeLo. After completing all calculations and scoring, PoMeLo prompts the user to provide the same NWK and CSV files of representative genomes (one per species) used in the genome size analysis. Once these files are uploaded, PoMeLo regenerates figures in the context of the phylogeny and adds trees to the visualizations. In cases of one-to-one comparisons between two species, the optional phylogenetic portion of the PoMeLo script may be skipped as it yields no useful information.

### Data Import

Once launched, PoMeLo automatically loads all required R packages and creates a new directory called `pomelo_outputs` to store all intermediate and output files. The user can change the name and location of the output directory according to their preference. PoMeLo presents a pop-up window prompting the user to select their “target genomes” file containing the list of pathogen genomes to be targeted for host-directed therapeutics. These should include genomes of the pathogen of interest and optionally genomes of similarly streamlined relatives. After the file is selected, PoMeLo accesses BV-BRC [10] and downloads the full pathway annotations for each genome individually. The same process is then repeated for the “non-target genomes” file containing the list of genomes of non-pathogenic or free-living organisms to be used for comparison with the target genomes of interest.

This process of downloading genomic annotations for target and non-target genomes is the most time-consuming part of the PoMeLo pipeline. Because the computational time increases linearly with the number of selected genomes, it is advisable to limit the total to fewer than 1000 individual genomes per analysis. Selecting one or a few representatives of each species for the analysis from the BV-BRC database will also help to optimize computational efficiency and reduce download time. Once genome annotation files have been downloaded to the working directory, PoMeLo combines all the data from both target and non-target genomes into a single comprehensive data frame. PoMeLo then prompts the user to find the “`mapping_GO_to_ecgene_and_ecpathway_toPATRIC.tab`” file that contains a list of the annotated metabolic pathways and corresponding genes in the BV-BRC database, allowing for mapping to a complete reference. PoMeLo integrates this information into the larger data frame to allow the identification of missing pathways and genes in the target and non-target genomes. PoMeLo also accesses BV-BRC and downloads the genome summary file to extract genome statistics for all genomes included in the analysis.

### Data Pruning

Before conducting further calculations, PoMeLo carries out a series of validation and name correction steps. First, the genome names are modified for simplicity and clarity, e.g. by abbreviating phrases like “`endosymbiont of …`”, removing terms like “`Candidatus`” and “`uncultured`”, and adding underscores after terms like “`sp.,`” “`strain,`” or genome project acronyms such as “`ATCC,`” “`FDAARGOS,`” “`NCTC,`” and “`USDA`”. Next, PoMeLo creates new columns that combine the first two names from both the “`Genome Name`” and “`Species`” fields for downstream merging of genome annotations with identical taxonomy. PoMeLo compares the “`Genome Name`” and “`Species`” fields and defaults to the name used in the “`Species`” field that has been manually curated by BVBRC. Furthermore, PoMeLo incorporates manual updates for recent changes to the KEGG pathway list. It excludes six pathways (KEGG pathways: p72, p231, p471, p472, p473, p1058) that are not included in BV-BRC annotations, and it adds annotations for four pathways (KEGG pathways: p220, p270, p470, p541) when their constitutive genes are present.

### Statistical Calculations

In the integrated data frame described earlier, each row corresponds to a genomegene pair. The dataset contains detailed information about the genome, gene, and associated pathway. Using this data along with the universal mapping file, PoMeLo calculates summary statistics at the individual gene level and the pathway level. These statistics describe the extent to which genes are identified within a given species or group, at both the individual gene and pathway level. Note that some genes can be assigned to multiple pathways, leading to ambiguity about their importance in any given pathway. Furthermore, the presence of these genes can artificially inflate the importance of included but irrelevant pathways (e.g. photosynthesis). To address this, three additional metrics are provided: the “promiscuity index”, which represents the number of unique pathways in which a specific gene is found, the average promiscuity index across each pathway, and the percentage of “promiscuous” genes in each pathway. These metrics help to refine the interpretation of gene and pathway significance.

### Percentage Differential Metric

To assess differential gene loss between groups within each pathway, we first calculate the number of genes identified in a given pathway for the target group and the non-target group, respectively. These counts are then divided by the total number of pathway genes identified in the target and non-target genomes combined (“`bvbrc_genes_in_pathway`”), which yields the percentage of the pathway that is missing in either the target or non-target group (“`perc_missing_in_target_group_bypathway`”, “`perc_missing_in_nontarget_ bypathway`”). We chose to use the “`bvbrc_genes_in pathway`” metric in lieu of the full number of possible genes in the pathway (as determined by the mapping file) because both the target and non-target genomes commonly undergo significant genome reduction. If we were to divide the count metrics by the full number of pathway genes, both groups would be missing a significant percentage of the pathway, making detection of significant gaps between the groups more difficult.

After calculating the percent missing in each group, these values are subtracted from each other to yield the differential in percentages between the groups. This provides a measure of the distance between the groups based on gene presence at the overall pathway level. In cases of genome reduction in the target group, “`perc_differential_non-target_bypathway`” will be high. Conversely, in cases where the target group has acquired additional genes via horizontal gene transfer or other mechanisms, the “`perc_differential_target_bypathway`” will be high instead. Although beyond the primary scope of PoMeLo, these measures could be utilized to identify areas in the target genomes that may have gained metabolic potential, providing insights into their function and pathogenicity.

### Calculation of Total Differences at Gene Level

While the most obvious signs of metabolic loss between groups may be observed in differences at the pathway level, it is also important to examine their gene-level differences. For example, while both target and non-target groups may encode 50% of the genes in the pathway, those individual genes might not overlap. In this case, the percent differential metric described above would not capture the differences between the groups, though informative data still exists to separate them. To address this, the algorithm calculates the percentage of genomes within each group that contain a specific gene, recorded as “`genepercentage_group_by_ecnumber`”. This value from the target group is then subtracted from that of the non-target group to yield the percent differential between groups for each gene, recorded as “`differential_bygene`”. To focus on pathway loss in the target group, all negative values are converted to zero. The “`differential_bygene`” values for each gene in the pathway are summed together and reported as “`total_differences_target_bygene`”.

### Predicted Metabolic Loss Score

To determine which pathways have undergone the most extensive reduction, we developed a scoring algorithm for Predicted Metabolic Loss (PML). This algorithm assigns an overall score to each pathway analyzed across the target and/or non-target genomes. The PML score is a positive integer calculated on a linear scale that is equivalent to the “`total_differences_target_bygene`” metric described above for each pathway. High PML scores indicate a greater degree of metabolic loss in the target genomes compared to the non-target genomes. The magnitude of the predicted metabolic loss score will vary between analyses, as the metrics depend on the genomes queried.

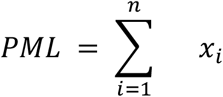

The PML score calculation is shown above, where *x* = “differential_by_gene”, *i* = a given gene, and *n* = the total number of genes in the pathway.

### Data Outputs

PoMeLo produces both tabular and visual output files in the ‘`pomelo_outputs`’ folder. The four tabular files are: 1) ‘`ec_number_pathway_stats`’, a catalog of individual genes and their corresponding pathways; 2) ‘`pathway_stats`’, a compendium of every gene-genome combination; 3) ‘`summary_of_ranked_pathways`’, which shows statistics by pathway; and 4) ‘`PML_fulldata_bypathway`’, a multi-tabbed spreadsheet of individual gene statistics for every pathway, ordered by the pathways’ PML score (Supplementary Tables 3-6).

PoMeLo generates visualizations in two different formats. In the first, single heatmaps summarize pathway presence (along the x-axis) by group, genus, or species (Supplementary Figures 2-4). In the second, multiple heatmaps – one per pathway – summarize gene presence and thus PML scores (along the y-axis) by group, genus, or species (Supplementary Figures 5-6). Running the phylogenetic steps will generate additional files, including phylogenies in both PNG & NWK formats (Supplementary Figure 7, Supplementary File 1), a figure combining phylogeny and genome size, and a final visualization combining phylogeny with pathway heatmaps, ordered by phylogeny (Supplementary Figures 8-9). To avoid an overabundance of outputs, only a selected subset of visualizations is saved in the ‘`pomelo_outputs`’ folder, while additional visualizations are saved in subfolders. All visualizations are generated in both PDF and PNG format, and each filename is appended with the date and a taxon name. The PoMeLo workflow is summarized in Figure 1.

**Figure 1.**
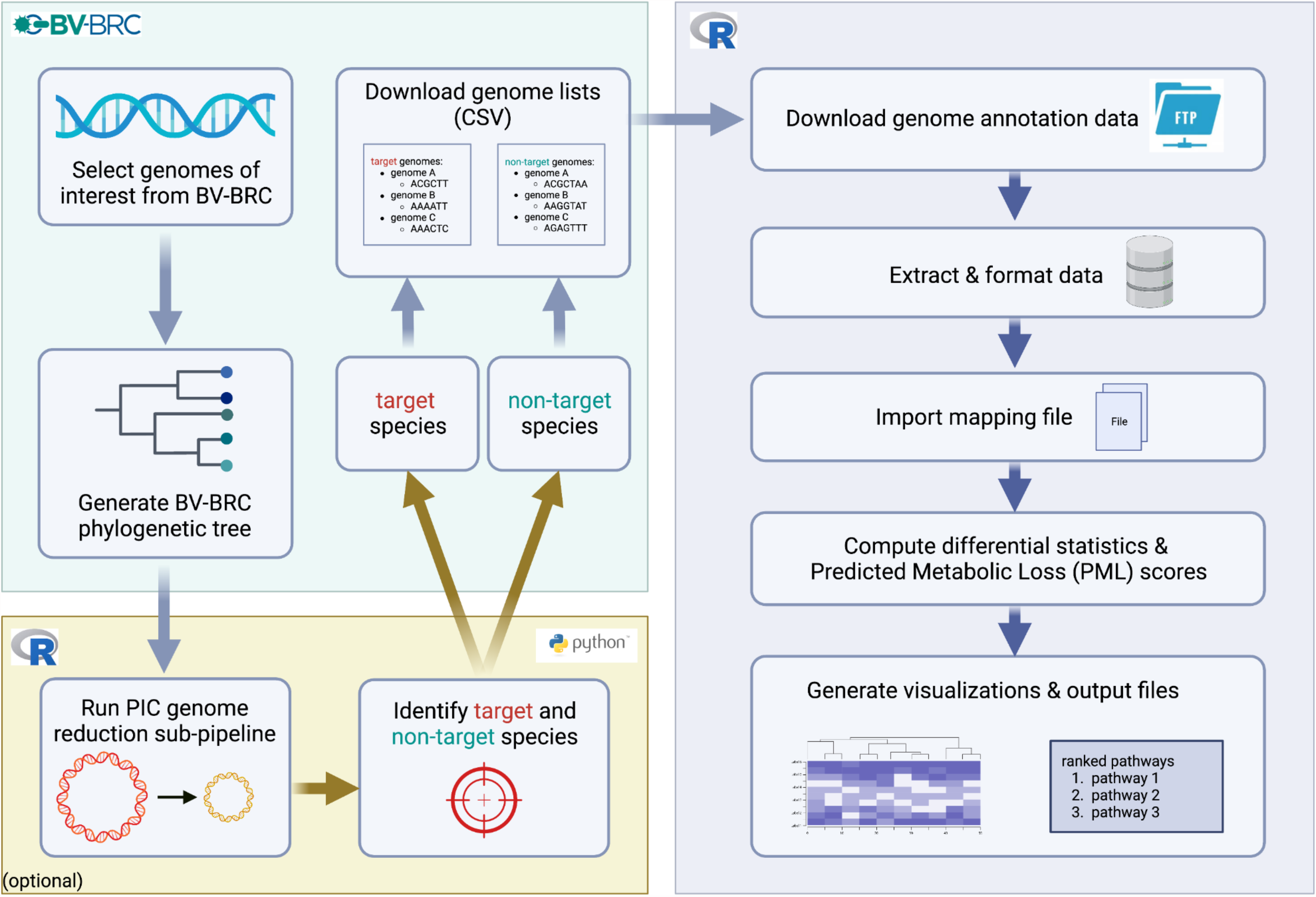
PoMeLo workflow.

### Computational Time

The time and computational requirements to run PoMeLo were assessed by conducting a comparative analysis with varying numbers and sizes of genomes included. Genome size can differ significantly across different taxonomic groups, from under 1 Mb to over 5 Mb, especially in the non-target group where major genome reduction is not expected. To account for this, we calculated computational time for different genome size ranges (<1Mb, 1-2Mb, 2-3Mb, 3-4Mb, and 4-5Mb). Because the number of genomes available through BV-BRC and included in the analysis can also vary significantly depending on the taxonomic group, we also varied this as a factor in our comparison with levels of 10, 50, 100, and 500 genomes.

Under all of these conditions, the comparisons were completed within 15 minutes of computational time when running PoMeLo locally on a Macbook Pro (5 GHz, M1, 8-core CPU, 8-core GPU, 16-core Neural Engine, 16 GB memory; Supplementary Table 7). The effect of genome size was minimal. This can be attributed to the fact that PoMeLo leverages pre-computed genome annotations generated by BV-BRC, rather than performing genome annotations *de novo*. Conversely, the number of total genomes in the analysis had a large effect on computational time, varying from ~3 min (2:37-3:24) for 10 genomes to ~13 minutes (11:18-14:20) for 500 genomes. Although computational run time is not a limitation, the authors nonetheless suggest limiting the number of genomes used to <500, to make interpretation of the results more manageable. It is important to note that the optional phylogenetic portion of PoMeLo is dependent upon the BV-BRC phylogenetic tree-building resource, which has a limit of 200 genomes. Users can avoid this limit by selecting a single representative genome for each species in the full analysis when building the phylogenetic tree. When running PoMeLo without the optional phylogenetic portion, there is no computational limitation to the number of genomes analyzed.

## Results

As a demonstration, we used PoMeLo to analyze genomes within the genus *Treponema*. Using the phylogenetically-weighted mean genome size analysis, we identified two species, *T. pallidum* and *T. paraluiscuniculi*, within a branch that is estimated to have undergone a genome size reduction of 49.6% compared to the estimated common ancestor (Supplementary Figure 10). *T. pallidum* and *T. paraluiscuniculi* have an average genome size of 1.1 Mb, in comparison to their closest relatives *T. parvum* (2.6-2.7 Mb), *T. vincentii* (2.6-3.0 Mb), *T. denticola* (2.7-3.1 Mb) and *T. phagedenis* (2.9-3.8 Mb). Therefore, we chose *T. pallidum* and *T. paraluiscuniculi* as the target group, with *T. phagedenis*, their closest evolutionary relative, as the non-target group. Using BV-BRC, we selected 52 genomes (51 of *T. pallidum*, and the 1 available genome of *T. paralusicuniculi*) in the target genomes file and 10 genomes of *T. phagedenis* in the non-target genomes file.

The PML scores for this comparison ranged from 0 to 2100, with a median score of 100 (Figure 2A). The target group encoded zero genes in 106/166 total pathways. For 37 of these pathways, the non-target genomes encoded at least one gene. The three pathways with the highest PML scores were purine metabolism (PML = 2100), amino sugar and nucleotide sugar metabolism (PML = 1500), and cysteine and methionine metabolism (PML = 1200) (Figure 2B). In these pathways, 12 to 21 genes were lost in the target species, representing a range of 58–75% of the genes encoding metabolic enzymes observed across all three species in the analysis. Because PML scores are a sum of the gene-level differences across a given pathway, pathways that have a small number of genes may have low PML scores, even in cases where the difference in pathway completeness is significant. Taken together, our results clearly indicate that the genomes of *T. pallidum and T. paraluiscuniculi* have become streamlined, and these pathogens have lost the ability to biosynthesize and metabolize multiple amino acids, carbohydrates, and nucleic acids in favor of relying on the host for these metabolites. As described in an accompanying manuscript (Medicielo et al.), this information can be used to identify known inhibitors of the missing metabolic enzymes as potential host-directed antimicrobials.

**Figure 2.**
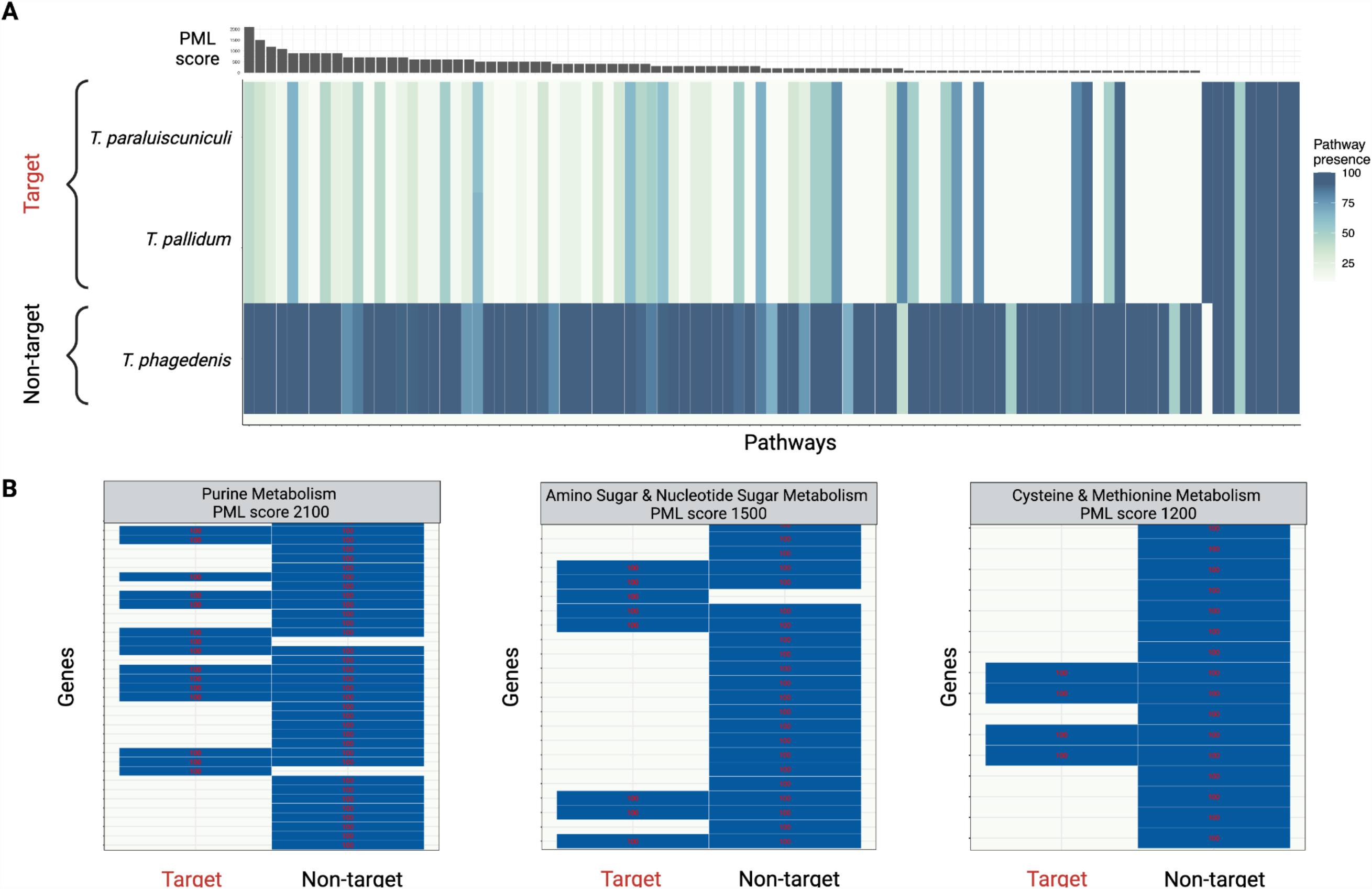
PoMeLo results in *Treponema*. The target group contains genomes of *T. pallidum* and *T. paraluiscuniculi*. The non-target group contains *T. phagedenis* genomes. (A) Completeness for all pathways in *Treponema*, shown at the species level. Color corresponds to percent pathway completeness; associated PML scores shown above the heatmap. (B) Pathways with the top 3 PML scores. Color corresponds to presence of individual genes in the group.

## Discussion

PoMeLo is a novel evolutionary genomics-guided computational approach to identifying metabolic gaps in pathogenic organisms using publicly available genomic data. The tool runs locally using user-friendly prompts and completes in a matter of minutes, with the largest determinant of computational time being the total number of genomes included in the analysis. PoMeLo generates both tabular and visual outputs that facilitate the comparison of predicted metabolic functionality between groups of genomes and individual species. It also algorithmically assigns each pathway a Predicted Metabolic Loss (PML) score to allow identification of genome streamlining at the pathway level.

Commonly used computational approaches for comparative metabolic profiling rely on large databases such as KEGG and RAST [7, 8, 14]. While these databases offer large pre-computed functional annotations for thousands of high-quality genomes, a major drawback of these approaches is that comparison across genomes is limited. PoMeLo leverages the existing structure of BV-BRC [10] to maintain the advantages of a centralized high-quality database of genomes, and it builds upon existing approaches by allowing the user to compare an unlimited number of genomes across individual genes as well as pathways. Furthermore, PoMeLo places the findings in an evolutionary context, which is critical for understanding the genome streamlining observed in pathogen genomes.

It is important to note that the accuracy and reliability of PoMeLo analyses depend on the quality of the genomes and associated annotations provided. This fact underlies the choice of utilizing BV-BRC for genome selection, which allows the user to screen their potential genomes for high quality and completeness. Expansion into lower quality genomes may be considered when the number of available genomes is limited, or when one complete genome is used along with several other lower quality genomes for validation with additional data points. The standardized annotation framework used by BV-BRC ensures better comparability across these genomes and annotations than those generated from different sources.

One potentially valuable application of PoMeLo is in the development of host-directed therapeutics. When pathogens lose the capability to generate certain metabolites, they pivot to siphoning nutrients from the host. By therapeutically inhibiting the production of these metabolites in the host, the pathogen can be starved of essential nutrients, aiding the host in clearing the infection [15–17]. Using PoMeLo to identify promising targets would reduce the time and cost associated with host-directed drug screening by repurposing existing drugs developed for other human diseases. It is important to validate PoMeLo findings *in vitro*, as we demonstrate in our accompanying manuscript where we (1) identify pathways to target for host-directed therapeutics using PoMeLo, (2) validate the targets *in vitro* in pathogen-infected cells, and (3) show using metabolomics that the PoMeLo-identified pathways largely overlap with pathways that are depleted in infected versus mock-infected cells, signaling pathogen metabolite stealing (Medicielo et al).

PoMeLo is currently only applicable for comparative analysis of bacterial genomes. Future directions could involve expanding its application to eukaryotic pathogens, for which evidence of genomic streamlining is also well-documented [18–20]. However, significant hurdles including the lack of high-quality and/or complete eukaryotic pathogen genomes, increased genome size and complexity, and the effects of polyploidy must be overcome. Other applications of PoMeLo may be to characterize the metabolic capabilities of entire microbial communities and the contributions of individual species to the overall metabolic profile. The extensive collection of genomes and annotations available at BV-BRC facilitates such an application. Combining PoMeLo analysis with metabolic profiling of the microbiome could be used to predict the effects of host-directed therapeutics on the host microbiome at various body sites[21].

In summary, PoMeLo is a novel computational approach for identifying metabolic gaps in genomes of pathogenic organisms. The tool represents an advance over existing methods by facilitating the comparison of an unlimited number of genomes and placing the results in evolutionary context. Through the analysis of genome streamlining, PoMeLo enables prediction of essential genes or pathways that could be targeted by host-directed therapeutics. PoMeLo is freely available for academic and non-academic use at https://github.com/czbiohub-sf/pomelo.

## Availability and requirements

All scripts, datasets and outputs described in this paper are available at: https://github.com/czbiohub-sf/pomelo. Project name: PoMeLo. Project home page: https://github.com/czbiohub-sf/pomelo. Operating system(s): platform-independent. Programming languages: R, Python. Other requirements: R 4.1.1 or higher, Python 3.8 or higher, R packages: `igraph`, `RColorBrewer`, `hexbin`, `scales`, `grid`, `lattice`, `gdata`, `gridExtra`, `ape`,, `reshape2`, `ggplot2`, `seqinr`, `phangorn`, `fs`, `hash`, `ggdendro`, `phytools`, `openxlsx`, `coop`, `tidyverse`, `rstuodioapi, aplot`, `BiocManager`, `BiocManager`(`remotes`), `BiocManager`(`YuLab-SMU/treedataverse`). License: none. Restrictions for non-academic use: none

## List of Abbreviations

BV-BRC: Bacterial and Viral Bioinformatics Resource Center
CSV: Comma-separated Values
KEGG: Kyoto Encyclopedia of Genes and Genomes
Mb: Megabases
NWK: Newick format
PDF: Portable Document Format
PIC: Phylogenetically Independent Contrasts
PML: Predicted Metabolic Loss
PoMeLo: **P**redictor **o**f **Me**tabolic **Lo**ss
PNG: Portable Network Graphics
RAST: Rapid Annotations using Subsystems Technology

## Declarations

### Ethics approval and consent to participate

Not Applicable.

### Consent for publication

Not Applicable.

### Availability of data and materials

All scripts, datasets and outputs described in this paper are available at: https://github.com/czbiohub-sf/pomelo, and we have used Zenodo to assign a DOI to the repository: 10.5281/zenodo.8172742

## Competing interests

The authors declare that they have no competing interests.

## Funding

This work was funded by the non-profit research institution Chan-Zuckerberg Biohub San Francisco in San Francisco, California.

## Authors’ contributions

A.G. & E.W. developed the tool, created and maintained the github repository and wrote the manuscript. G.D. developed the code for the phylogenetically independent contrasts analysis, provided guidance on topics related to evolutionary biology and edited the manuscript. J.W. lead the computational biology team developing the tool, provided feedback and edited the manuscript. V.A. developed the concept for the tool, directed the project and edited the manuscript.

## Acknowledgements

Josette Medicielo provided guidance and feedback throughout the development of the tool and tested the tool. Sandy Schmid provided feedback and guidance on the tool and the manuscript.

## Notes

https://github.com/czbiohub-sf/pomelo

### Competing Interest Statement

The authors have declared no competing interest.

