## Supplemental Figures for "PoMeLo: a systematic computational approach to predicting metabolic loss in pathogen genomes"

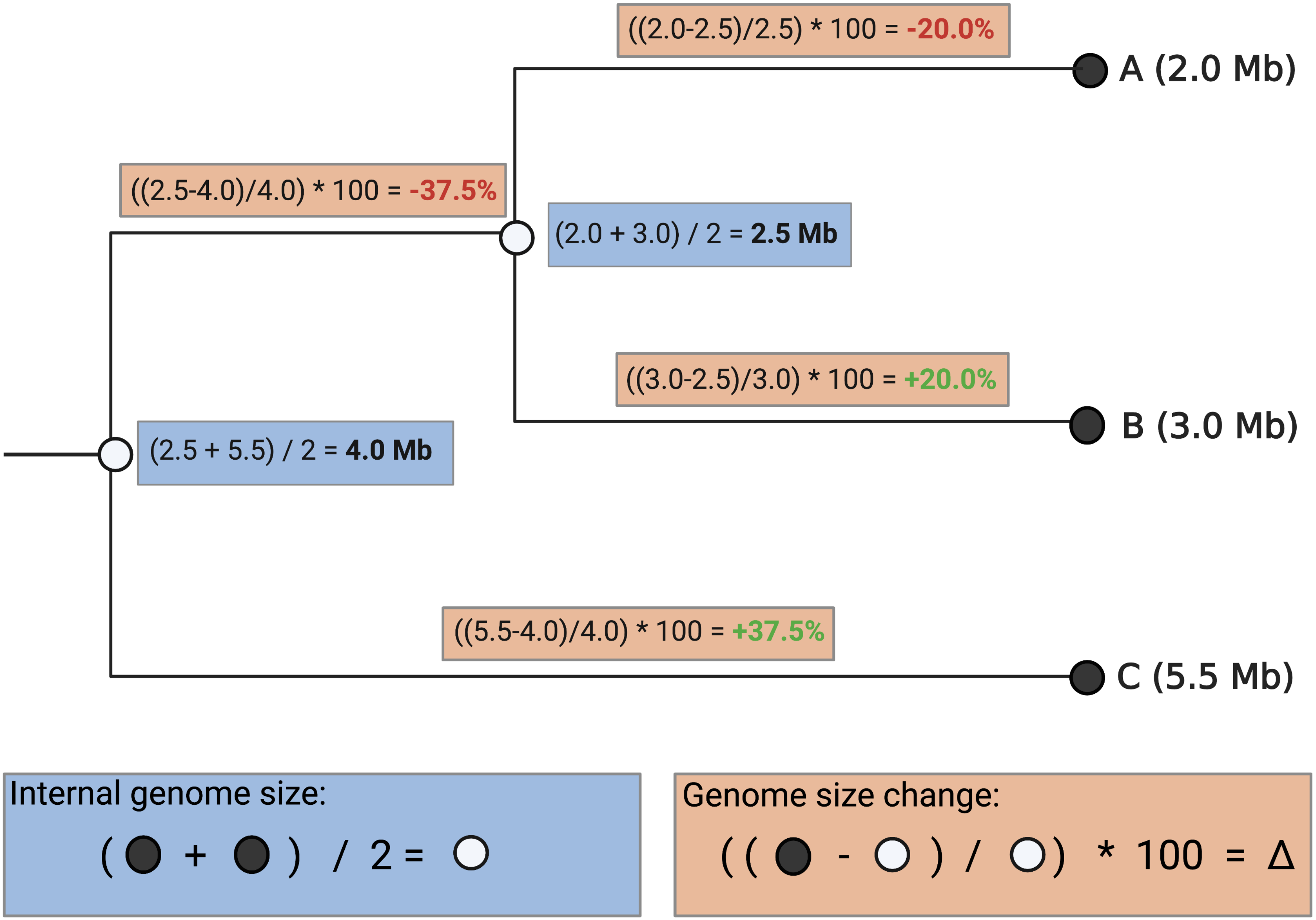


Supplementary Figure 1. Schematic of phylogenetically weighted mean genome size calculations. Black nodes are labeled A, B, & C with known genome size in megabases (Mb) in parentheses. Blue boxes contain genome size calculations for internal white nodes. Orange boxes along the branches contain calculations of genome size changes between nodes (red = loss; green = gain) .

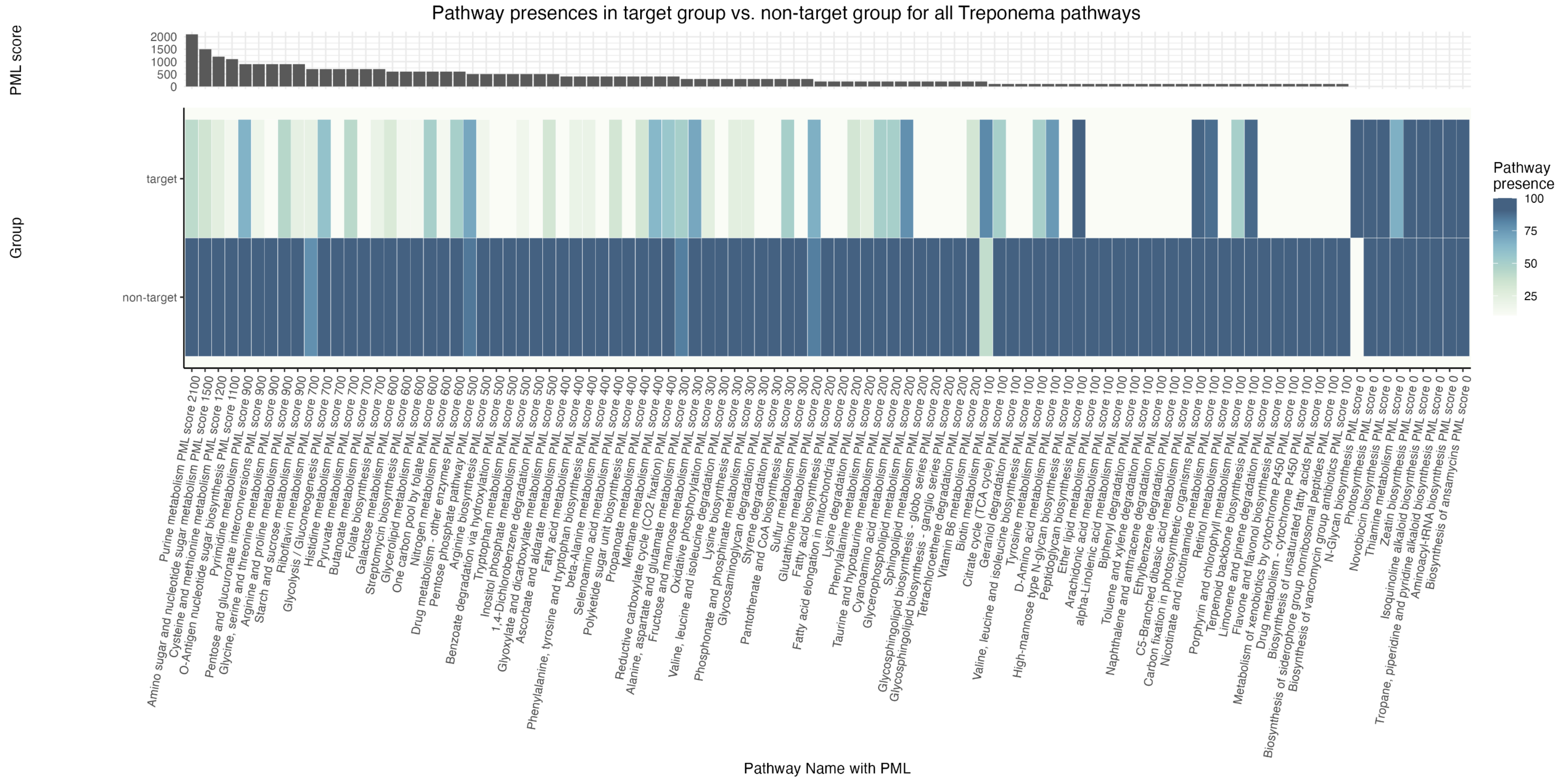


Supplementary Figure 2. Heatmap of all pathways, by group. Pathways are shown on the x-axis and the target and non-target groups are shown on the y-axis. Pathways are ranked by PML score , descending  left to right. PML scores are shown at the top of the plot. The color of cells corresponds to the percentage of genes identified in a given pathway for a given group.

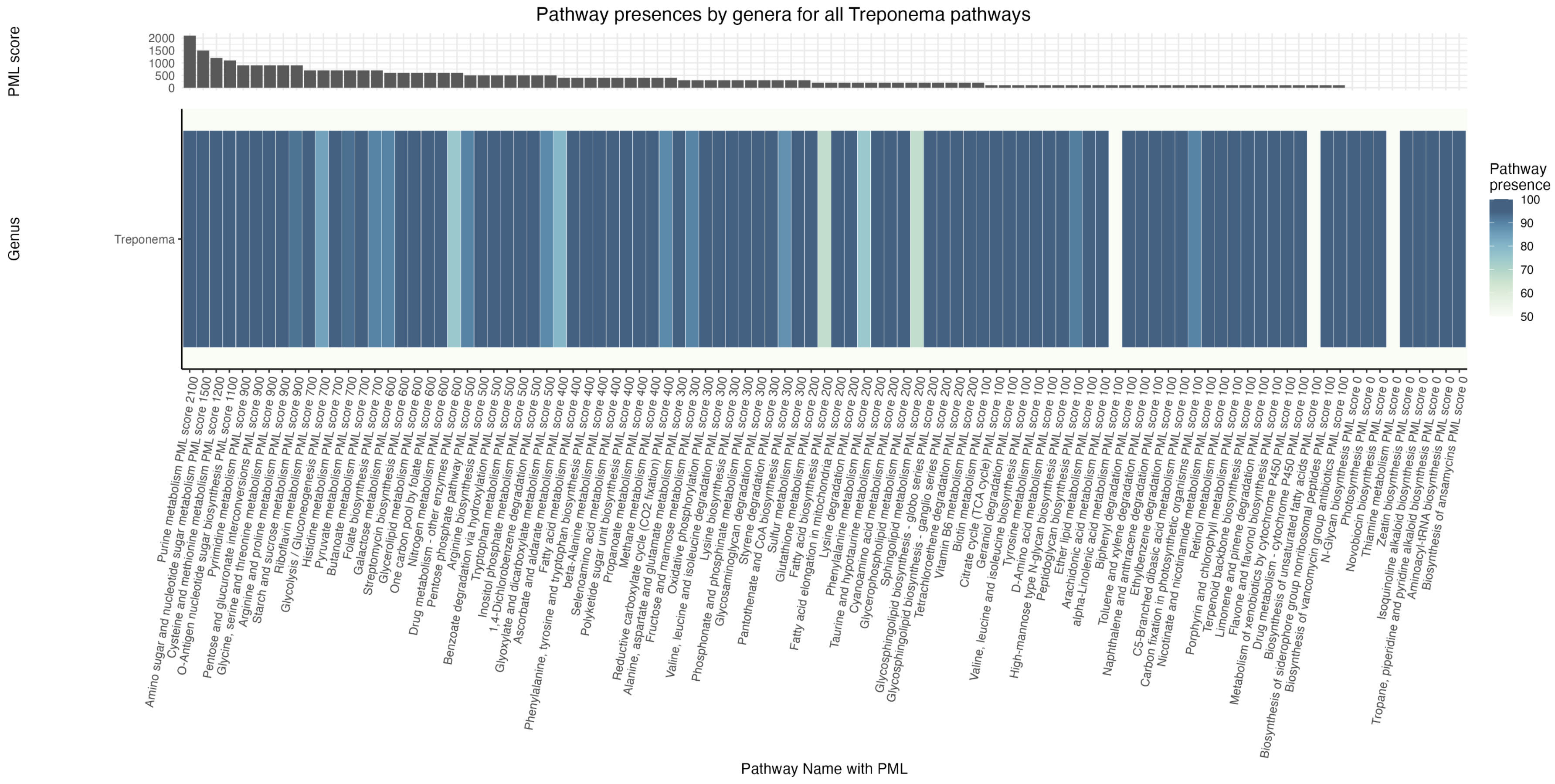


Supplementary Figure 3. Heatmap of all pathways, by genus. Pathways are shown on the x-axis and genera are shown on the y-axis. Pathways are ranked by PML score, descending from left to right. PML scores are shown at the top of the plot. The color of cells corresponds to the percentage of genes identified in a given pathway for a given gen.

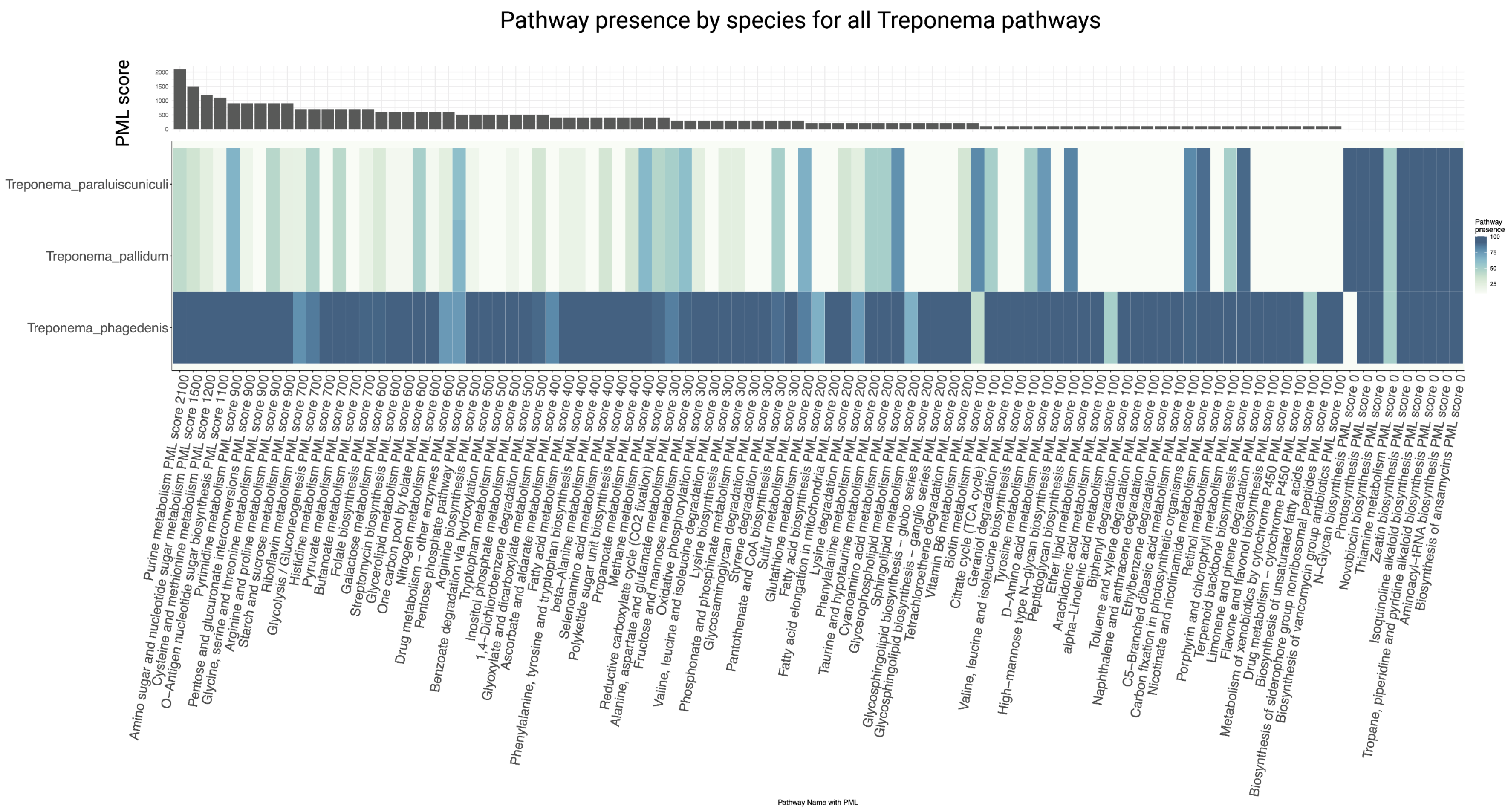


Supplementary Figure 4. Heatmap of all pathways, by species. Pathways are shown on the x-axis and species are shown on the y-axis. Pathways are ranked by PML score , going from high to low and left to right. PML scores are shown at the top of the plot. The color of cells corresponds to the percentage of genes identified in a given pathway for a given species.

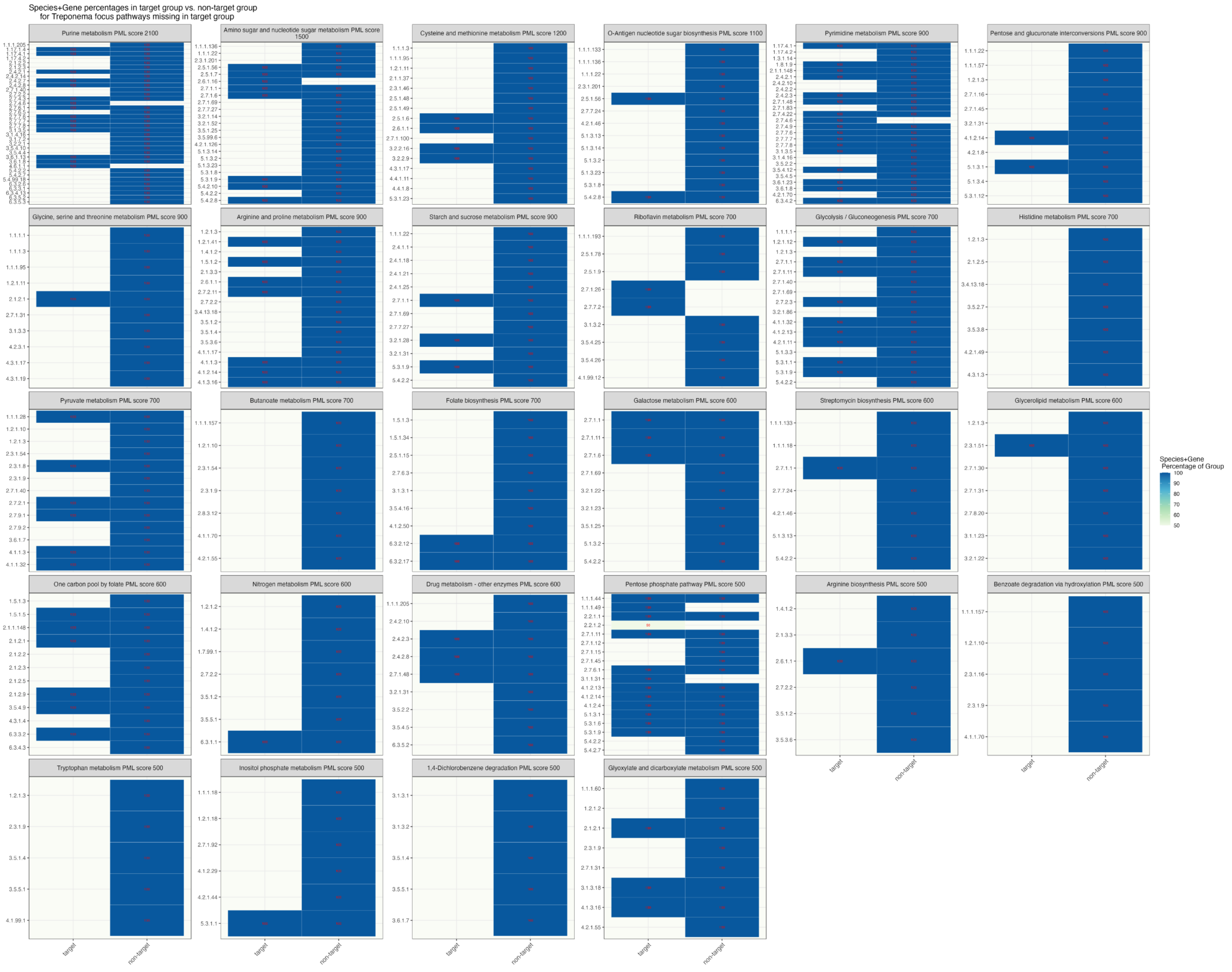


Supplementary Figure 5. Heatmaps of high PML pathways, by group. Each pathway is represented by its own heatmap. Groups (target & non-target) are shown on the x-axis and individual genes, here shown as Enzyme Commission (EC) numbers are shown on the the y-axis. Heatmaps are ordered by highest PML score to lowest, from left to right and top to bottom. Pathway names and PML scores are shown above the heatmaps. The color of cells corresponds to the percentage of genomes within a given group for which a given gene was identified. In cases where cells are blank, zero genomes in the given group were found to encode for this particular gene. The percentage of genomes encoding the gene in a given group is also shown in red text within the corresponding cell.

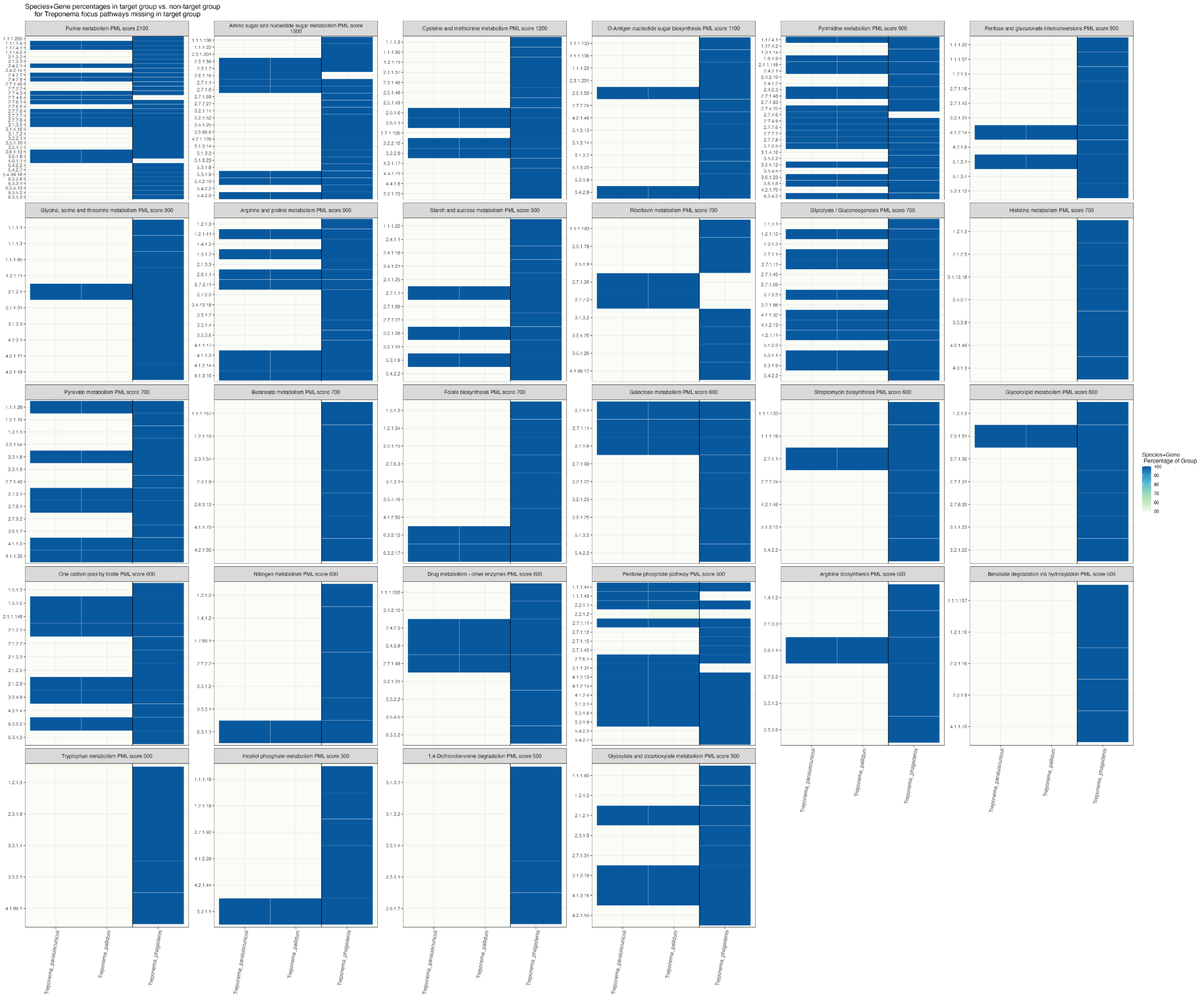

Supplementary Figure 6. Heatmaps of high PML pathways, by species. Each pathway is represented by its own heatmap.Species are shown on the x-axis, separated by group, and individual genes, here shown as Enzyme Commission (EC) numbers are shown on the  the y-axis. Heatmaps are ordered by highest PML score to lowest, from left to right and top to bottom. Pathway names and PML scores are shown above the heatmaps.  The color of cells corresponds to the percentage of genomes within  a given species for which a given gene was identified. In cases where cells are blank, zero genomes in the given species were found to encode for this particular gene. The percentage of genomes encoding the gene in a given species is also shown in red text within the corresponding cell.

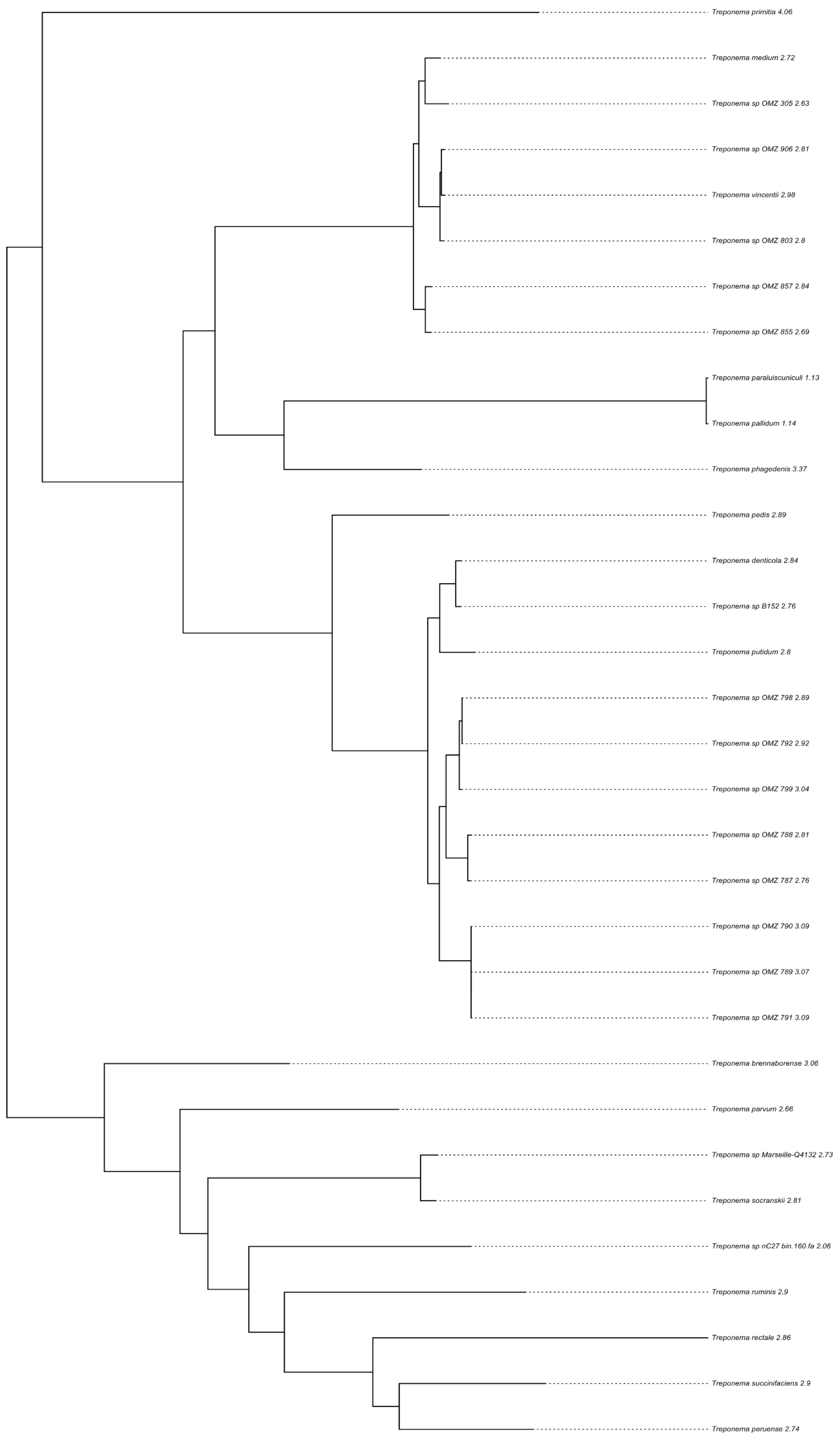


Supplementary Figure 7. Treponema phylogeny. Phylogenetic reconstruction of Treponema genomes generated by PoMeLo, based on BV-BRC generated NWK file and genomes included in analysis.

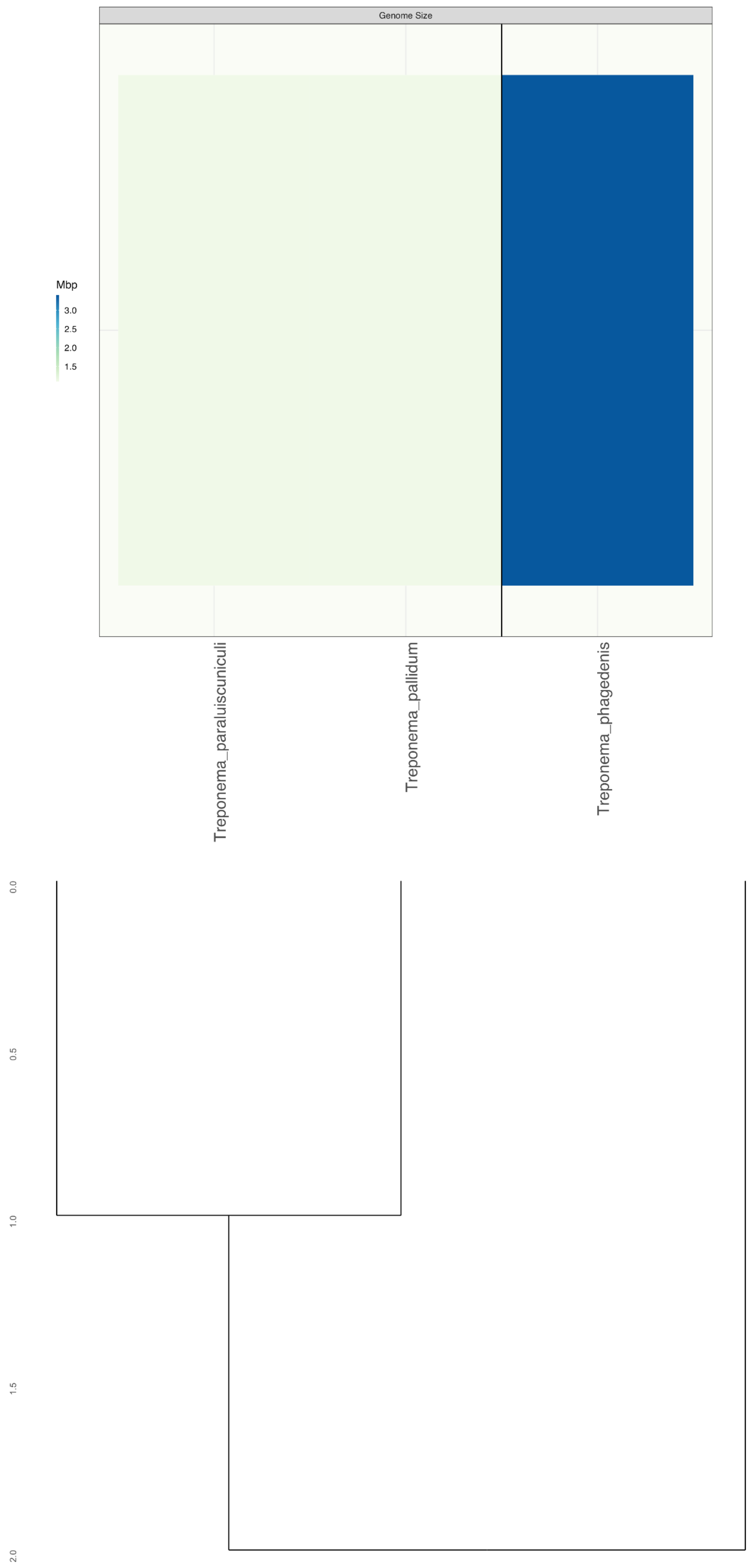


Supplementary Figure 8. Treponema Genome Size Phylogeny. Phylogeny generated by PoMeLo based on Bv-BRC generated NWK file and genomes in the analysis. Phylogeny of Treponema species shown, separated by group (target vs. non-target). Color of cells corresponds to genome size in megabases.

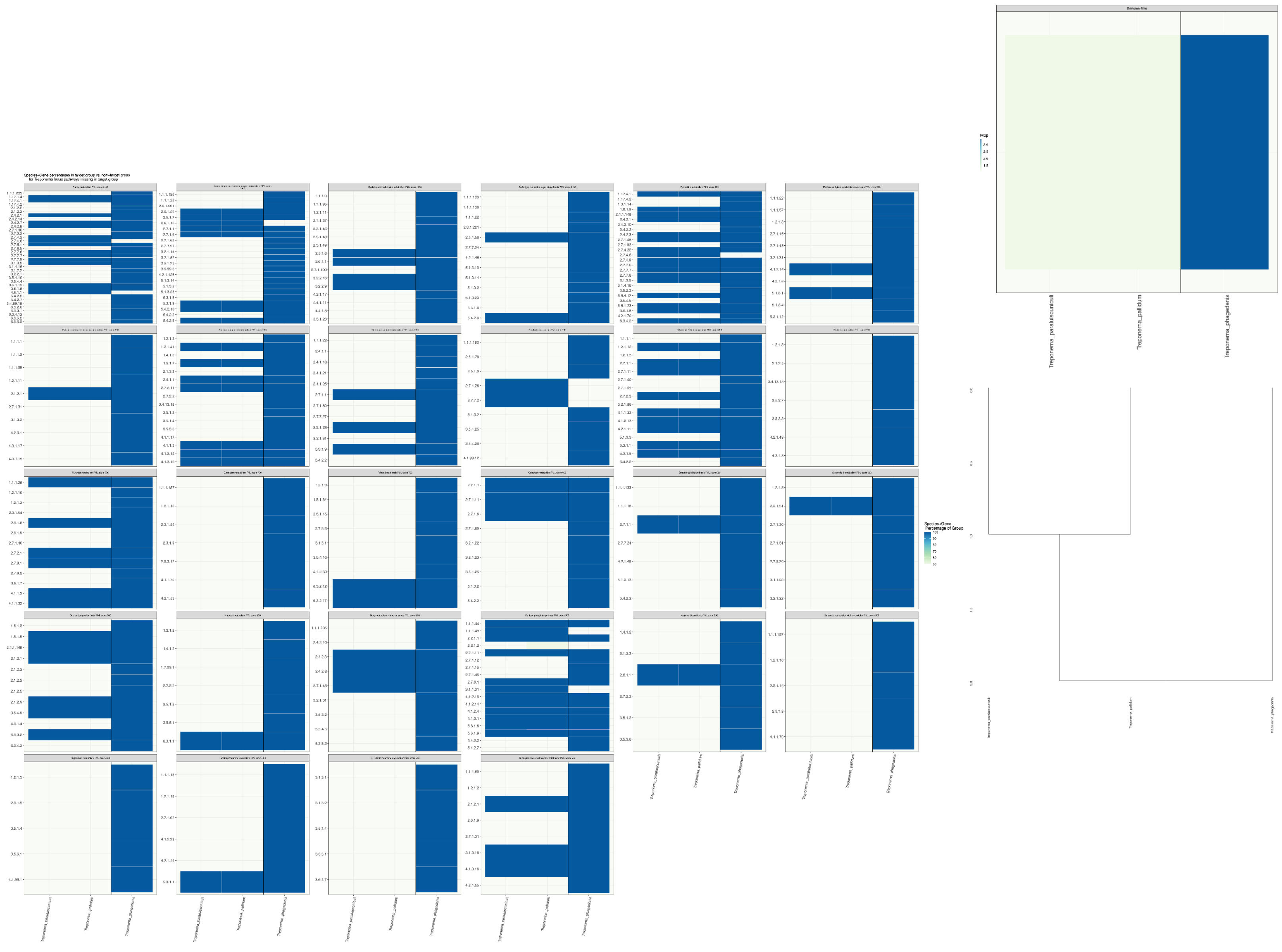


Supplementary Figure 9. Heatmap of high PML pathways ordered by phylogeny.  Each pathway is represented by its own heatmap.Species are shown on the x-axis, separated by group.Individual genes, shown as Enzyme Commission (EC) numbers are on the  the y-axis. Heatmaps are ordered by highest PML score to lowest, from left to right and top to bottom. Pathway names and PML scores are shown above the heatmaps.  The color of cells corresponds to the percentage of genomes within a given species for which a given gene was identified. In cases where cells are blank, zero genomes in the given species were found to encode for this particular gene. A phyolgenetic tree of all species is shown to the right of the heatmaps. Colors of cells correspond to the genome size in megabases.

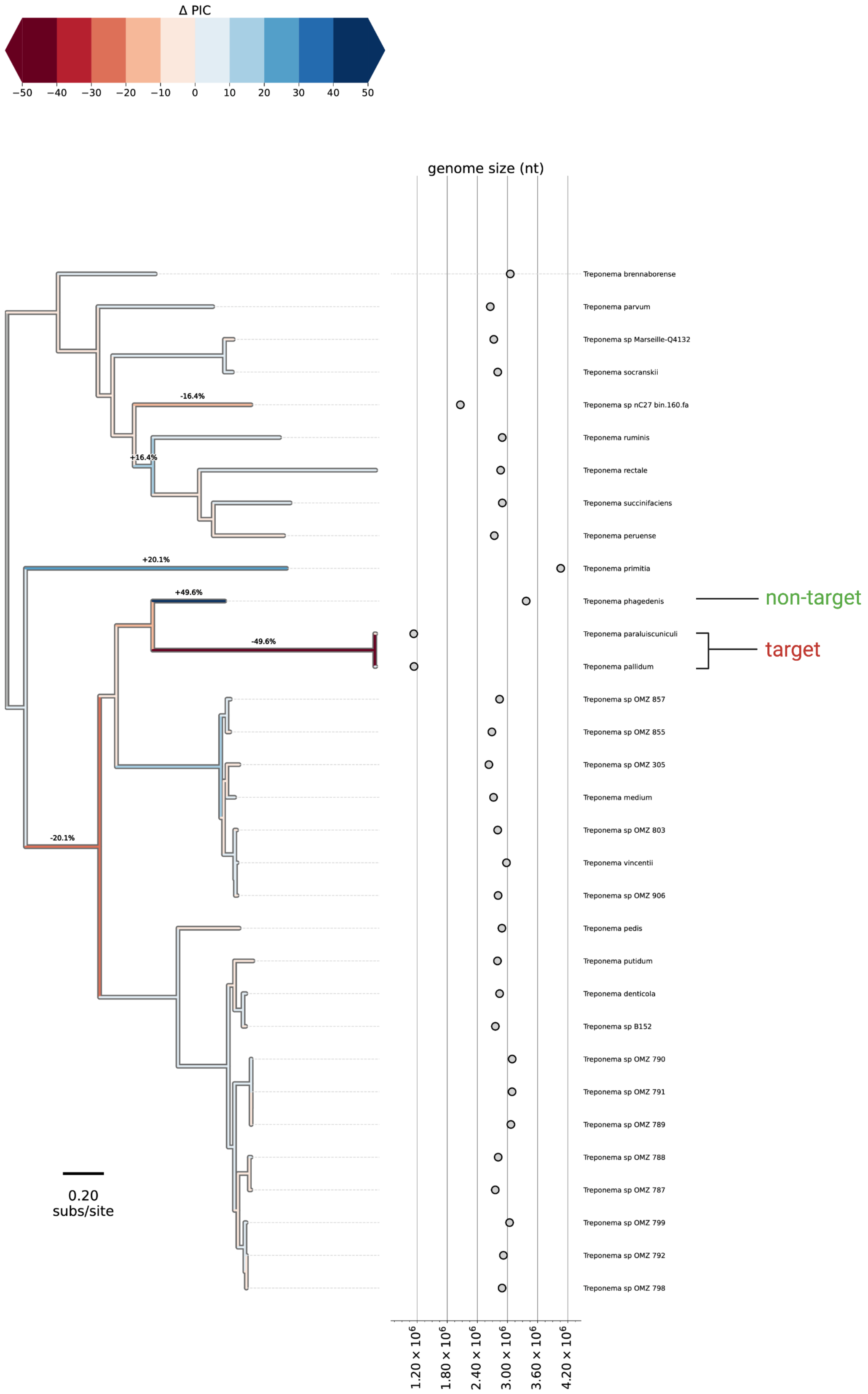


Supplementary Figure 10. Phylogenetically-weighted mean genome size analysis of *Treponema pallidum*/*Treponema paraluiscuniculi* and related organisms. All genomes downloaded from BV-BRC. Length of branches represents the evolutionary distance between nodes based on substitutions per site. Color of branches corresponds to the change in genome size in comparison to the common ancestor. Genome size in nucleotides is plotted on the right. Changed in genome size of less than -10% or greater than 10% are shown.
