## Supplemental Table 7 for "PoMeLo: a systematic computational approach to predicting metabolic loss in pathogen genomes"

| | Total Number of Genomes | | | |
|---|---|---|---|---|
| **Genome Size (Mb)** | **10** | **50** | **100** | **500** |
| **<1 Mb** | 2:37 | 3:20 | 4:07 | 11:18 |
| **1-2 Mb** | 2:53 | 3:48 | 4:35 | 12:53 |
| **2-3 Mb** | 3:05 | 3:50 | 4:42 | 13:01 |
| **3-4 Mb** | 3:07 | 4:10 | 5:00 | 13:32 |
| **4-5 Mb** | 3:24 | 4:12 | 5:08 | 14:20 |

Supplementary Table 7. Computational Time Analysis. All times shown in minutes:seconds (m:s) format.
